# Silver Armor Against Bacteria: A Battle of Antimicrobial Effectiveness

**DOI:** 10.1101/2024.08.03.606483

**Authors:** Alina Zaidi

## Abstract

For many farmers, there is a need to improve crop resistance to pathogenic organisms. Florida’s climate and hurricane-prone location promotes the spread of many crop pathogens making management difficult and expensive. Therefore, this study evaluates using a Do-It-Yourself, DIY, method to produce colloidal silver solutions that may be used as effective inhibitors of bacteria growth. The efficacy was compared across gram-negative and gram-positive bacteria species. Production of colloidal silver used an electric current from eight 9-volt batteries wired in series. The starting silver was a jewelry chain, approx. 98% silver, suspended in a container of distilled water. Treatment effects were compared to a commercially available silver solution (10K, ppm) as the positive control *via* the Kirby-Baur method. Each petri dish was divided into four quadrants, into which each had a treated cellulose square impregnated with a treatment solution. Treatments were: T1-10K ppm AgNp; T-2-5ppm AgNp, T3-3ppm AgNp; and positive control T4-blank water control. The dimensions of Zones of Clearance (ZOC) and the bacterial growth surrounding treated squares were analyzed. The experiment was replicated four times. Data analysis conducted using one-way ANOVA with *post hoc* separation of means using the T test and Tukey’s HSD. The positive control solution was the most effective bactericide across all species, followed by Treatment-2 (5-ppm treatment), which caused ZOC in gram-negative species. Treatment-3, the 3ppm, did not significantly affect bacterial suppression, while activity at 5ppm suggests that simple home-based, DIY systems can produce low cost, bactericidal nano-silver solutions. In this experiment the bacterium that tested that is also beneficial to plants, *R. rubrum*, showed an increased tolerance to all silver treatments. Improving homemade, DIY, systems may provide low-cost treatment solutions against some bacteria species important to backyard agriculturists.

## Introduction

Nanosilver inhibition of common agricultural pathogens, such as *Serratia marcescens*, provides insight into emerging disease prevention methods. Unlike common bactericidal treatments, silver nanoparticles (AgNP) enter through porin proteins while accumulating, and damage cell membranes (Jung et al, 2008; Tran et al, 2013; Ahmad, et 2020; More et al, 2023). Protein synthesis is often hindered *via* ribosomal decay, which lends to antibacterial properties (Ahmad et al., 2020; Vila Domínguez et al, 2020). Commercialized silver products are sold as health supplements, cleaning products, and are used to coat many household appliances (Sim et al., 2018). Silver’s properties can be manipulated using colloidal silver treatments, a solution which suspends silver particles in a solvent. The efficacy of bactericidal effects is linked to AgNP size, shape, and concentration (Guilger-Casagrande et al., 2019; More et al, 2023; Park et al, 2011; Tran et al, 2013). Smaller nanoparticles are more effective (Tran et al, 2013 Lu et al., 2013) to kill cells *via* DNA and RNA impairment (Rai et al., 2009; Tripathi and Goshisht 2022). In citrus trees, trunk injection leads to silver transportation throughout the xylem and phloem of the tree, producing a systemic bactericidal effect (Su et al., 2020). Gram-negative bacteria species are becoming more virulent in healthcare settings due to the growth of antibiotic resistance to modern healthcare products (Pereira et al., 2022), this includes *B. catarrhalis* (Catlin 1990). Gram-negative species contain an extra outer membrane which make them more resistant to a variety of antibiotics (Breijyeh et al., 2020). For example, *Serratia marcescens* (Class: Gammaproteobacteria) is a gram-negative pathogen which causes black rot in citrus and other agricultural disease (Hasan et. al, 2020). The bacterium, *S. marcescens* has been documented to have developed virulent antibiotic resistance (Roy et. al 2023). Approximately 75% of strains of *Branhamella catarrhalis* (Class: Gammaproteobacteria) produce β-lactamase and are therefore resistant to penicillin and penicillin derivatives (Nissinen et al, 1995). *Rhodosprillium rubrum* (Class: Alphaproteobacteria) uniquely benefits agriculture, leading to increased resistance to pathogenic strains of bacteria and increased cell viability (Madnay et al., 2022). The gram negative species tested were susceptible to the microbicidal effects of silver at concentrations greater than five ppm. This is due to destruction of the cell wall caused by biologically active metals (Breijyeh et al., 2020). Gram-positive species differ from those above, with a thicker cell wall. The gram-positive bacteria included: *Micrococcus luteus* (Fleming 1928) (Class: Actinomycetes), in the family Micrococcus, a Gram-positive cocci broadly found in natural environments such as soil and water resources and it is usually considered a normal inhabitant of human skin and oropharynx mucosa (Erbasan 2018). *Bacillus cereus* (Frankland 1887) (Class: Bacilli) is a gram-positive species widely spread in poultry and spice farm environments, identified as a food-borne toxin (Gill et al., 2024). In this study, homemade, or Do It Yourself, DIY, silver solutions harness may provide a cost-effective antibacterial for those looking for DIY methods. Researchers continue to demonstrate significant suppression of bacterial pathogens (Vila Domínguez et al., 2020).

## Materials and Methods

Five bacteria species were purchased as part of a BSL-1 kit from Carolina Biological Supply (Item 154615). Gram-negative cultures: *Branhamella catarrhalis, Rhodospirillum rubrum*, and *Serratia marcescens*. Gram-positive cultures: *Bacillus cereus* and *Micrococcus luteus*. The stock silver solution was provided by the US Department of Agriculture. Colloidal silver was produced using a system of a silver jewelry chain (9 Inches), and two quarters (1963). The method is described online at “How To Make Colloidal Silver” https://www.youtube.com/watch?v=-jSQqiOvifw). Treatments were: T1-10K ppm AgNp; T-2-5ppm AgNp, T3-3ppm AgNp; and positive control T4-blank water control. In brief required materials included: alligator clips, eight 9-volt batteries, glass jars, and distilled water (Pure Life, Zephyrhills, Florida).

### Colloidal Silver

The eight 9-volt batteries were attached to the silver component using wiring. The electrical current (72 volts) system was run for 24 and 48 hrs to allow silver particles to suspend in the water. A volt-meter (TDS Meter, iSpring) displayed silver concentrations at each time point measured.

### Kirby Baur Method

Using 60 nutrient agar Petri dishes in four separate trials, the Kirby-Baur Method was used to determine bacterial suppression across treatments. Each plate was streaked with one of the five bacterial species (*B. cereus, B. catarrhalis, M. luteus, R. rubrum, S marcescens)*. Squares (2×2) of sterilized paper (Profile, Louisville, Kentucky, 177°C) were impregnated with each treatment of nanosilver solution. Each petri dish was then divided into four quadrants and a treatment square was placed in each. The four treatments: Treatment-1, 10,000 ppm (positive control), Treatment-2, 5ppm, Treatment-3, 3ppm, and Treatment-4, distilled water (negative control). Each of the four treatments were placed in each plate. After 24, 48, and 72 hrs, the Zones of Clearance (ZOC) around each treatment square were measured to quantify bacterial suppression (Fig. 1).

**Figure 1.**
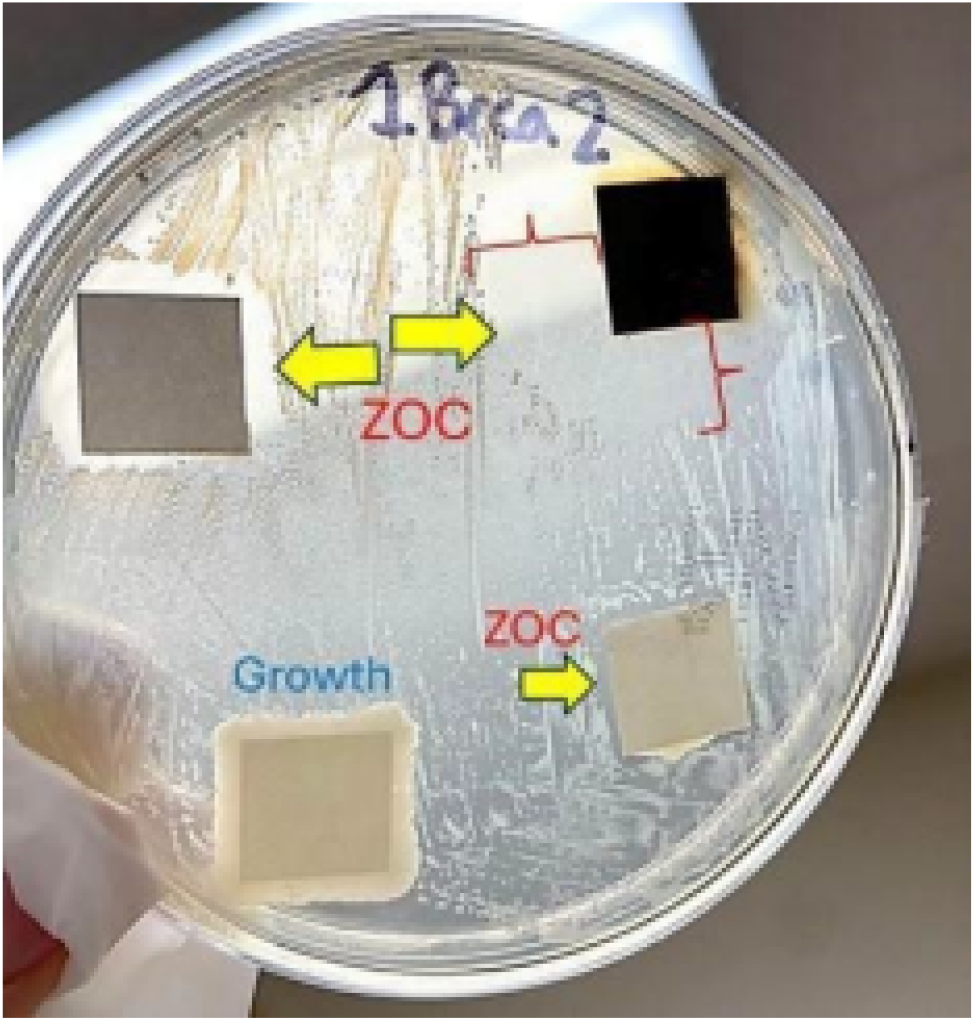
Example of Zones of Clearance, ZOC, 24 hrs after treatment for *Branhamella catarrhalis.*

### Bacterial Growth and ZOC Testing

Petri dishes were treated with bacterial species. The plates were then treated with all four silver treatments *via* the Kirby-Baur Method. The Petri dishes were incubated at 33°C for 24hrs. The ZOC around each treatment square was recorded in millimeters. Plates were incubated for another 24 hrs. Additional bacterial growth surrounding selective treatment squares was noted. The ZOC and Bacterial Growth were quantified using two measurements across the lateral diameters. Petri plates were incubated for another 24 hrs and results were recorded, at 24, 48, and 72 hrs (Fig.2).

**Figure 2.**
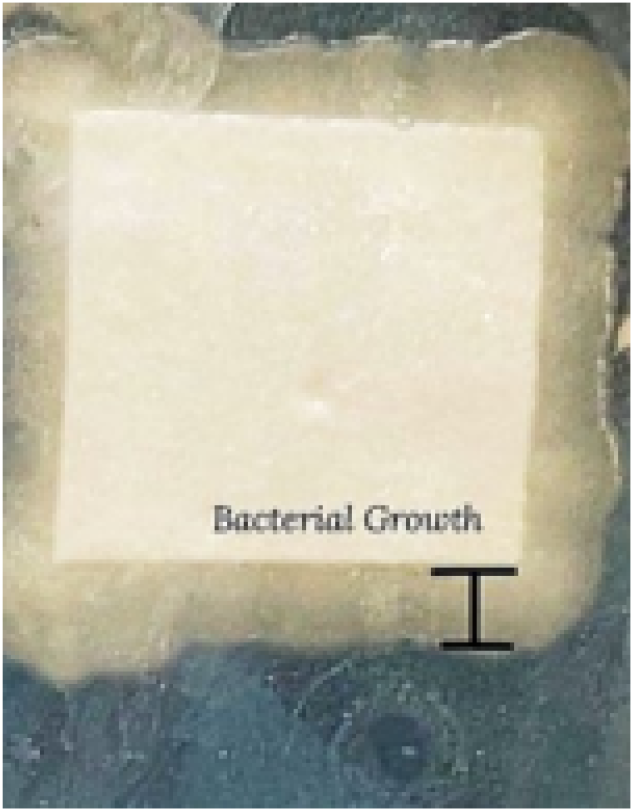
Example of Bacterial Growth surrounding AgNPs, 72 hrs after treatment for *Branhamella catarrhalis*.

### Statistical Analyses

The data were summarized as the mean ± SE (standard error) for all data sets. The data were then subjected to a One-way analysis of variance (ANOVA) (Lakens 2013) using Analyse-it® Statistical analysis add-in for Microsoft Excel, significance, Excel (version 7.2.10 68) and/or ANOVA using Social Science Statistics (https://www.socscistatistics.com/, accessed 18 August 2023). Means separation post-hoc Tukey’s Test with differences considered statistically significant at the 5% confidence level (p ≤ .05).

Statistical analyses were conducted using the website https://www.socscistatistics.com/. The One Way ANOVA for Independent Measures, with post hoc separation of means using Tukey’s Test, *P* < .05.

## Results and Discussion

This study quantified ZOC *via* the Kirby Bauer method but uniquely also recorded increased bacterial growth around the treatment squares after 72hrs of incubation. The data of this concentrated growth is referred to as Bacterial Growth. ANOVA’s one-way T-test was used to determine statistical significance and Tukey’s HSD was used to compare differences between treatment means.

### Zone of Clearance, ZOC – Figure 4

The positive control, Treatment-1 (10k ppm) was significant from all other treatments for all bacteria except the gram-negative, *R. rubrum*.

For *B. catarrhalis*, the ANOVA: F(3,80) = 53.3, p < .05, post-hoc comparison of means using Tukey’s Test (*P* ≤ .05). Treatment-2 had statistically significant differences in ZOC, 14.88 mm, with Treatment-3 (3.13 mm) and Treatment-4 with no ZOC (0.0 mm) (Fig. 3). *For S. marcescens*, ANOVA: F(3,80) = 235.1, p<0.001, *post-hoc* comparison of means using Tukey test (P ≤ .05). Only Treatment-1, 10k ppm had statistically significant (ZOC=16.37 mm) For *M. luteus*, ANOVA: F(3,80) = 14.9, p < .05, post-hoc comparison of means using Tukey’s Test (*P*≤ .05). The positive control, (10k ppm) the ZOC =18.50 mm, and in Treatment-2 (5ppm) the ZOC= 2.50 mm. While Treatment 3 (3 ppm) did not produce statistically significant difference (*p>0*.*1*, ZOC = 0.19 mm).

**Figure 3.**
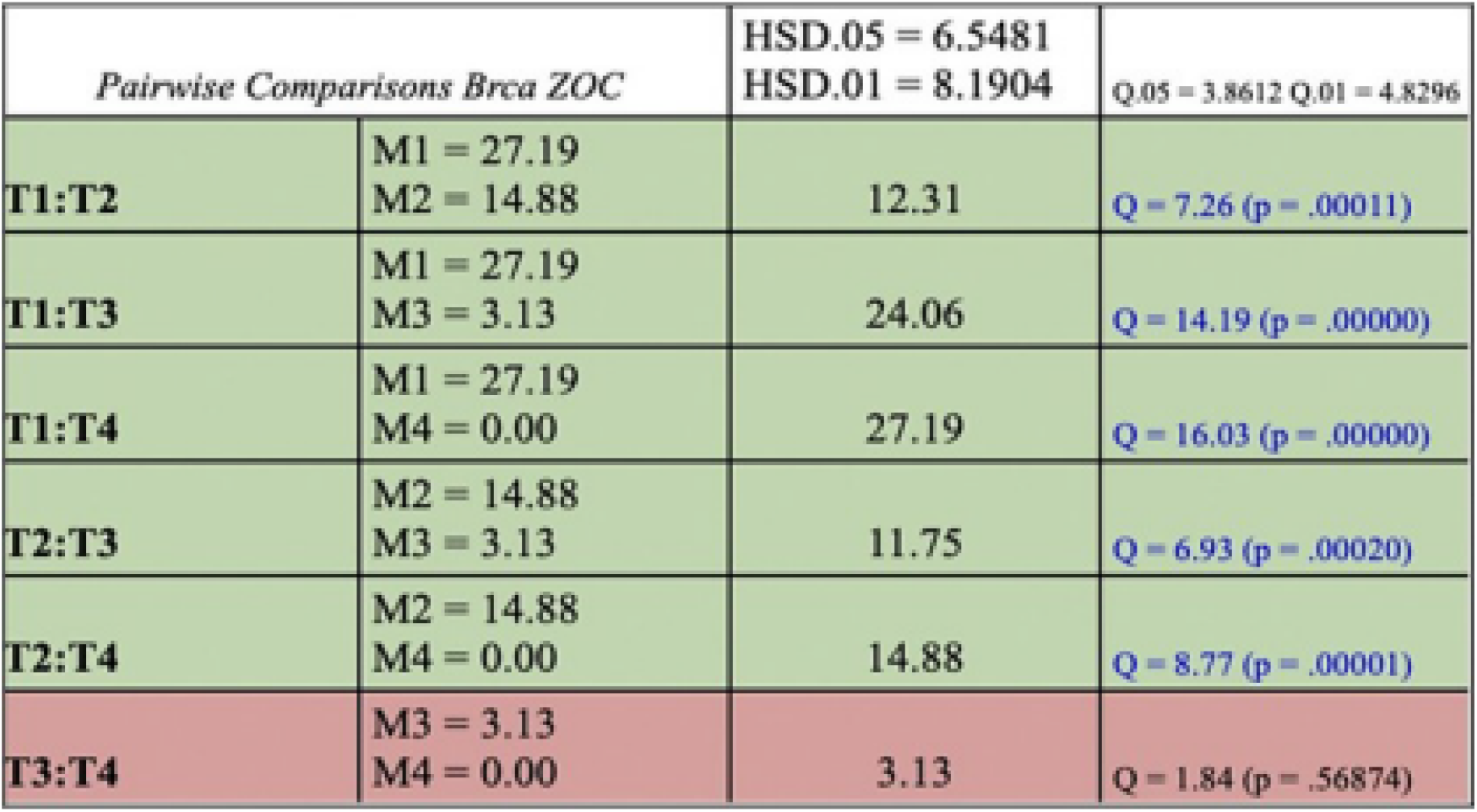
Zones of clearance at 24hrs for *Branhamella catarrhalis* compared to all other treatments.

**Figure 4.**
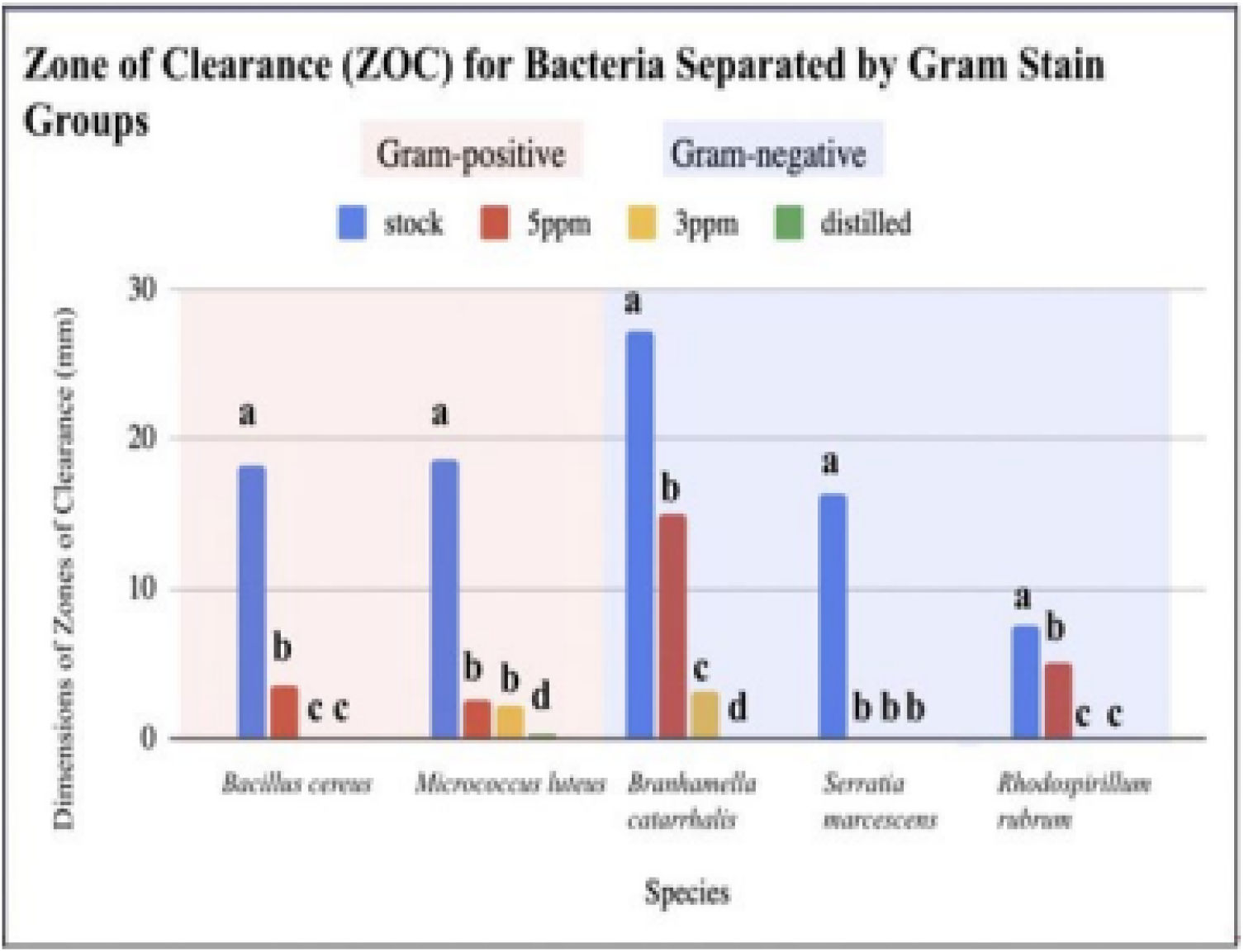
Zones of clearance, ZOC, at 24 hrs. Image separated as gram-positive bacteria (Pink background, *Left*) and gram-negative bacteria (Blue background, *Right*). Bars labeled with different letters within bacterial species show statistically significant difference at *P* < .05

For *B. cereus*, ANOVA: F(3,80) = 53.3, p < 0.001, post-hoc comparison of means using Tukey’s Test (*P* ≤ .05). Treatment-2 (ZOI =2.50 mm) was not statistically significantly different (p = 0.9). Treatment-3 (p =0.19331) had no statistically significant difference from the distilled water control (p = 0.9).

For *R. rubrum*, ANOVA: F(3,80) = 1.6, p < .05, *post-hoc* comparison of means using Tukey’s Test (*P* ≤ .05). were all not statistically significantly different from the water control treatment. Treatment-1 (p = 0.3), Treatment-2 (p= 0.5) and Treatment-3 (p < .05).

### Bacterial Growth -Figure 5

For *B. catarrhalis*, ANOVA: F(3,80) = 1.88, p = 0.2, post-hoc comparison of means using Tukey’s Test (*P* ≤ .05). There was no statistically significant difference between all treatments. For *S. marcescens*, ANOVA: F(3,80) = 3.23, p < .05, post-hoc comparison of means using Tukey test (*P* ≤ .05). Bacterial growth of Treatment-3 (11.38mm) was statistically significantly different from Treatment-1 (0.00 mm).

**Figure 5.**
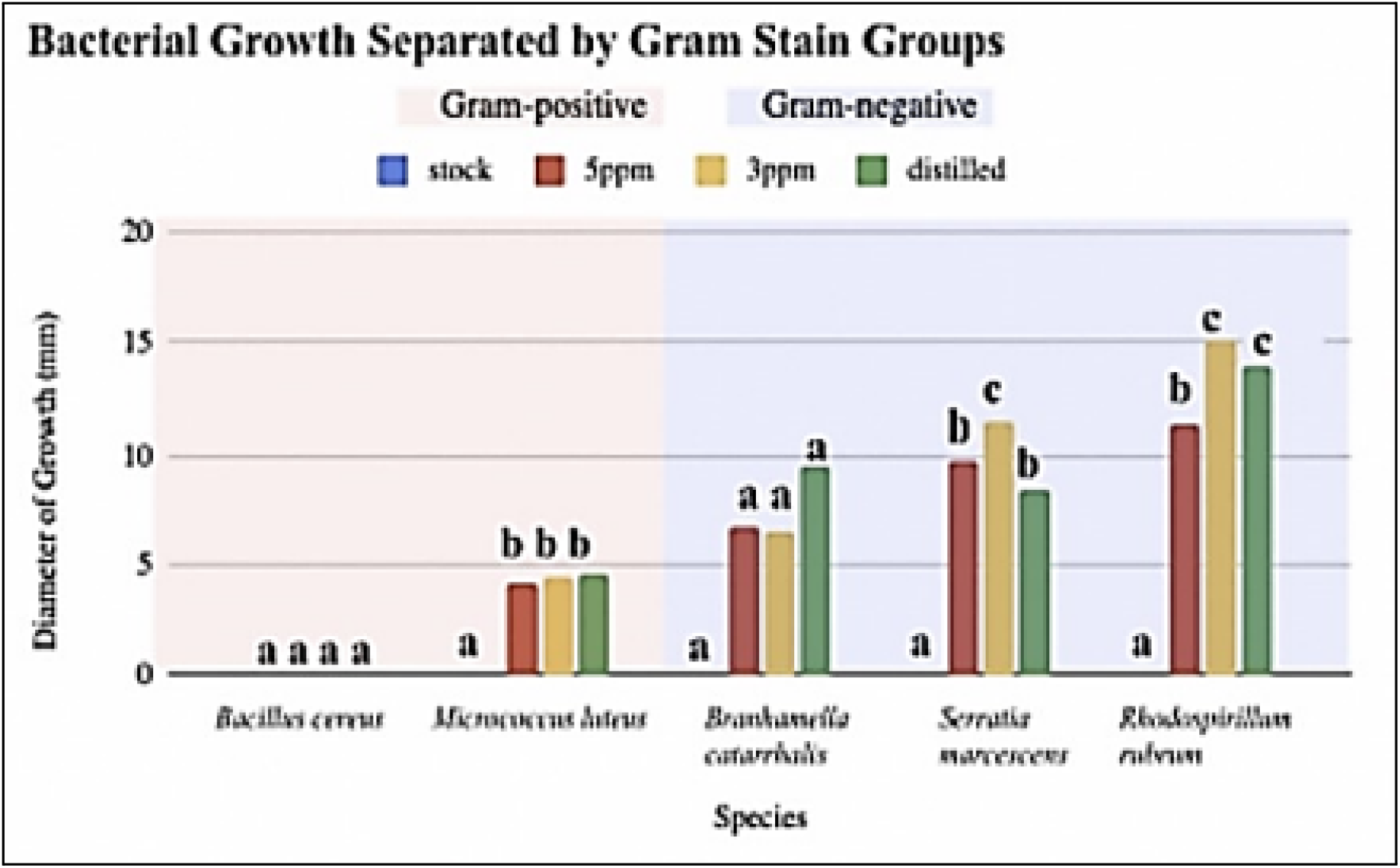
Effect of AgNp Treatments on Bacterial Growth at 72hrs. Data is separated as gram-positive bacteria (Pink background, *Left*) and gram-negative bacteria (Blue background, *Right*). Bars labeled with different letters within bacterial species show statistically significant difference of means at *P* < .05.*Treatment Effects on Gram-negative and Gram-positive Bacteria*

For *M. luteus*, ANOVA: F(3,80) = 0.78, p = 0.5, post-hoc comparison of means using Tukey’s Test (p < .05). There was no statistically significant difference between all treatments. Treatment-2 (4.19mm) (p = 0.6), Treatment-3 (4.44mm) (p = 0.6), and Treatment-4 (4.56mm) (p = 0.6).

For *B. cereus*, there was no statistically significant differences among all treatments. For *R. rubrum*, ANOVA: F(3,80) = 4.69, p < .05, post-hoc comparison of means using Tukey’s Test (*P* < .05). Statistically significant differences existed between Treatment-1 and Treatment-3 (15.00mm) (p < .05), and Treatment-1 and Treatment-4 (13.64 mm) (p < .05).

### Mechanistic Delivery on Gram-negative and Gram-positive Species

Figure 4 shows that colloidal silver solutions had more significant effects of bacterial suppression on the Gram-negative species. The outer membrane unique to gram-negative bacteria species is an effective defense mechanism against antibiotic agents. Gram-negative bacteria are susceptible to nanometal treatments due to damage to the outer membrane (**Fig. 6**) (Umair Raza et.al 2023)

**Figure 6.**
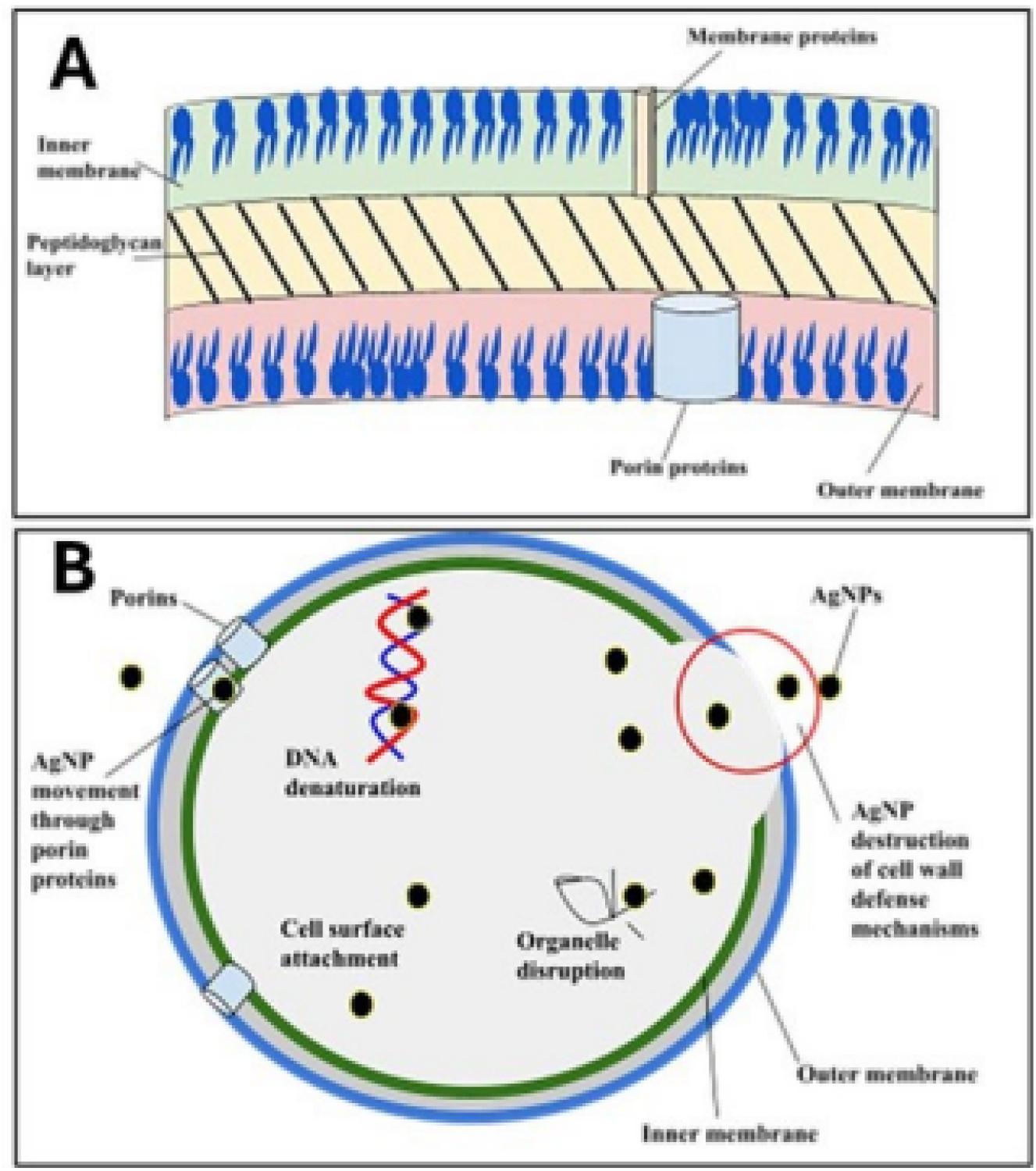
(A) Structures of gram-negative cell wall. (*image after*, Breijyeh et.al 2020). **(B) AgNPs impede outer Gram-negative bacterial defense mechanisms**. AgNPs attach themselves to the cell surface, enter through the cell wall, and deform the internal cell structures. (*image after*, More et.al 2023).

**Figure 7.**
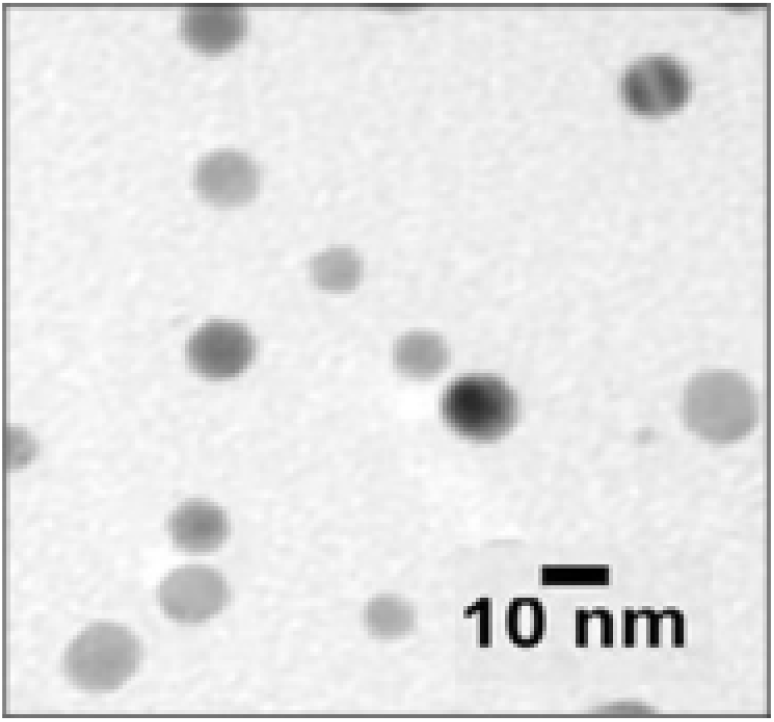
Average 10 nm diameter Silver Nanospheres, 10,000 ppm. Aqueous, 2mM Sodium Citrate. (nanoComposix 4878 Ronson Ct Ste J, San Diego, CA 92111. USA. Cat. No.: SKU: AGCN10-50M).

Bacteria are more susceptible to the antibacterial effects of smaller-sized AgNPs, 10 nm to 20 nm (Abbaszadegan et al, 2015; Agnihotri et al, 2014; Bao et al, 2015; Feng et al, 2000). Colloidal silver created with a greater voltage current results in smaller-sized particles and thus higher voltage systems should be implemented to improve DIY, homemade production to improve bactericidal activity.

### Stock Serial Dilution

The efficacy of a commercially available AgNP solution was directly compared to the concentrations of homemade, DIY, produced colloidal silver solutions using a serial dilution. ZOC and Bacterial Growth were recorded after 24, 48, and 74hrs for *B. catarrhali*s. Results show that the positive control AgNP solutions loses efficacy at concentrations below 100 ppm (**Fig. 8**.

**Figure 8.**
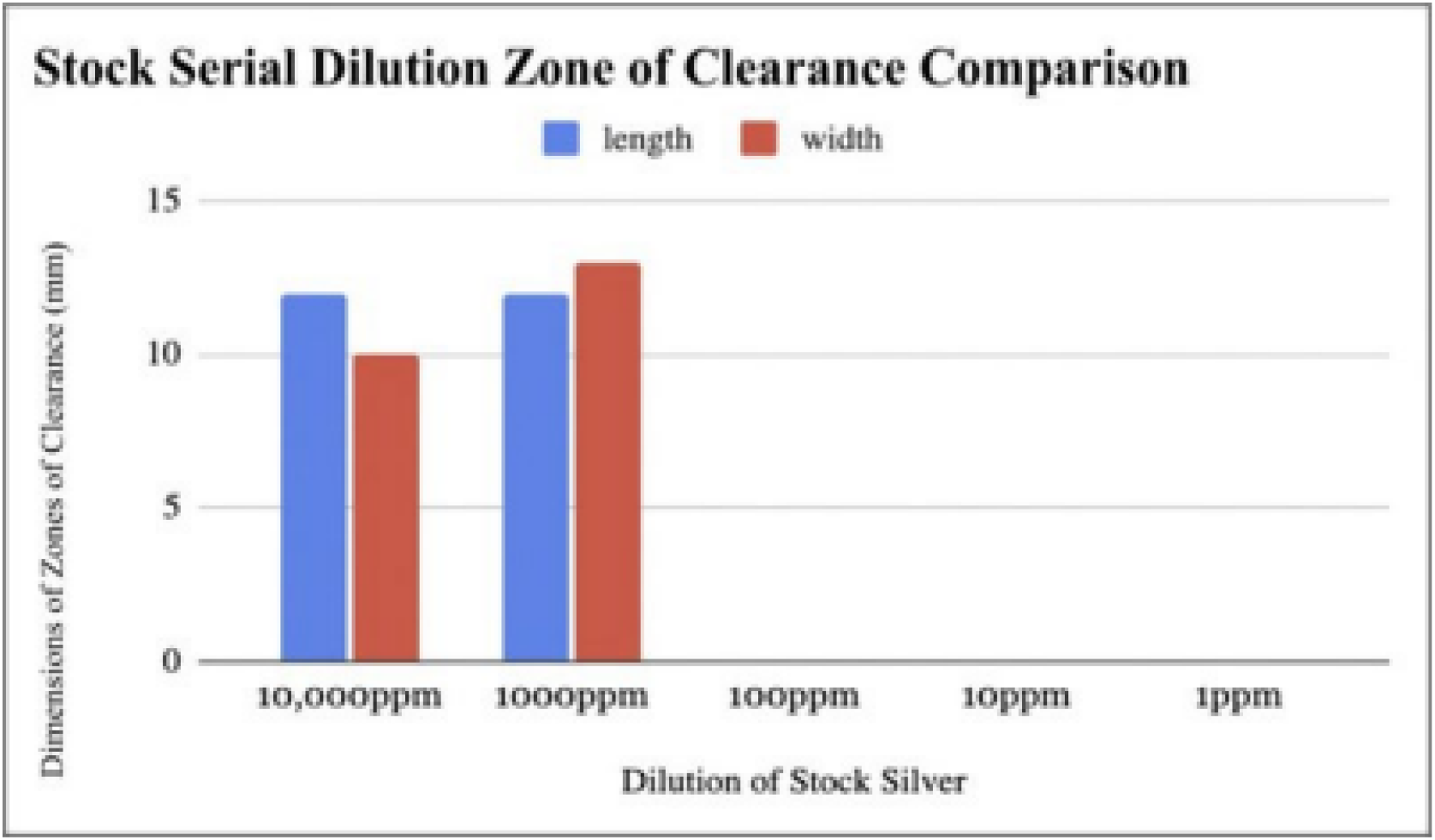
ZOC for *B. catarrhalis* after 24hrs. The ZOC produced by the commercial silver stock solution in serial dilution (1 ppm to 10,000 pm). Efficacy of bacteria suppression was observed at 1,000 ppm and 10,000 ppm, but not was not effective at concentrations below 100 ppm.

### Cost Analysis

A limited review of retail colloidal silver products compared to a homemade, DIY solution reveals that homemade colloidal silver can be approx. 350% lower in cost to produce, compared to the same concentration and quantity of commercial store-bought alternatives $25 – 45 USD.

## Conclusion

The greatest antibacterial effects occurred with the stock solution (Treatment-1, 10k ppm) which also had the smallest nanoparticles, 10 nm. Out of all the species evaluated, *B. catarrhalis* showed the greatest sensitivity to the nanosilver treatments. *Bacillus cereus* and *Micrococcus luteus* also showed significant sensitivity at the 5ppm treatment, more so than *Rhodospirillum rubrum* which did not exhibit sensitivity under all nanosilver concentrations.

The *R. rubrum* specie, which promotes plant growth demonstrated an inverse response to silver, with increasing growth as silver concentrations decreased. Some plants also respond positively to nanosilver treatments (Mehmood 2018; Hussain et al, 2019; This study demonstrates that the gram-negative bacterial species tested show a greater sensitivity to silver nanoparticle treatments supporting previous studies. The review by Breijyeh et al., (2020) also found that gram-negative bacteria are often more resistant to antibacterial products and antibiotics in healthcare settings. Research by Su et al, (2020) reported that silver nanoparticle treatments on citrus trees are effective against a gram-negative bacterium, *Candidatus* Liberibacter asiaticus. This research was further validated showing that nanosilver applied as a foliar spray also improved citrus fruit quality (Umair Raza et al, 2023).

Commercial products that claim to have the same concentrations of AgNP, yet produce different efficacies may be due to variations in the size, shape, or concentration of the AgNPs (Abbaszadegan et al, 2015; Feng, 2000; Lalegani et al, 2020; Tripathi and Goshist 2022; More et al, 2023). Some of the observed differences may also be the cause of unknown proprietary blends or compositions, such as surfactants, adjuvants, etc. These could also contribute to the antibacterial static, or antibacterial effects (Barillo et al, 2014; Leitão et al, 2018; Prema et al, 2022; Tran et al, 2013).

In conclusion, the results of my study demonstrate the activity of silver nanoparticle solutions on most of the bacteria tested. For pathogens, these solutions are already being commercialized for use. In healthcare settings, Sim et al (2018) finds that antimicrobial silver is expanding in terms of therapeutic treatments and bacterial resistance (Hossain 2021; Jung 2008; Sampath et al, 2022). For crops, silver applications are very recently being explored. A study by Umair Raza et.al (2023) found that for new citrus pathogens, silver solutions can be developed as effective disease management method. This research supports that silver could be used to develop silver treatment solutions as bactericides for use in crops. Moreover, exploring how to improve the production of colloidal silver solutions with smaller diameter AgNp using homemade, or DIY systems may contribute a valuable part of this emerging technology.

## Acknowledgements

I wish to thank, Dr. Wayne Hunte, USDA, ARS, U.S. Horticultural Research Laboratory, Fort Pierce, FL for mentoring and guidance on experimental design and statistics.

## Disclaimer

Mention of trade names or commercial products herein is solely for the purpose of providing specific information and does not imply recommendation or endorsement, to the exclusion of other similar products or services by the U.S. Department of Agriculture. USDA is an equal opportunity provider and employer.

## WEB-links

“How To Make Colloidal Silver.” YouTube, uploaded by School of Permaculture, 10 August 2016, https://www.youtube.com/watch?v=-jSQqiOvifw.

For ANOVA, T-test, and Tukey’s HSD: Glen, S. 2014. Chi Square P Value Excel: Easy Steps, Video. From StatisticsHowTo.com: Elementary Statistics for the rest of us! https://www.statisticshowto.com/probabilityand-statistics/excel-statistics/calculate-chi-square-

## SDS Citations

1. Silver MSDS sheet - https://www.sigmaaldrich.com/US/pt/sds/aldrich/576832
2. Educational Bacteria site; Source of Organism: Carolina Labs - https://www.homesciencetools.com/product/gram-stain-bacteria-comparison-set/3.
3. Bacterial Handling Guidelines - https://www.cdc.gov/orr/infographics/biosafety.html4.
4. Antiseptic Guidelines - https://www.tmcc.edu/microbiology-resource-center/labprotocols/aseptic-technique
5. Antiseptic Technique - https://blink.ucsd.edu/safety/research-lab/biosafety/containment/bsl1.html
6. MSDS Sheet From Supplier for Specific Bacterial handling: https://drive.google.com/file/d/1c_cIjXnOTf8a9Pn4HOilNOt33SldJOeq/view

